# GWAS identifies novel risk locus for erectile dysfunction and implicates hypothalamic neurobiology and diabetes in etiology

**DOI:** 10.1101/283002

**Authors:** Jonas Bovijn, Leigh Jackson, Jenny Censin, Chia-Yen Chen, Triin Laisk-Podar, Samantha Laber, Teresa Ferreira, Craig Glastonbury, Jordan Smoller, Jamie W Harrison, Katherine S Ruth, Robin N Beaumont, Samuel E Jones, Jessica Tyrrell, Andrew R Wood, Michael N Weedon, Reedik Mägi, Benjamin Neale, Cecilia M Lindgren, Anna Murray, Michael V Holmes

**Affiliations:** Big Data Institute, Nuffield Department of Medicine, University of Oxford, Oxford OX3 7LF, UK; Institute of Biomedical and Clinical Science, University of Exeter Medical School, University of Exeter, Exeter, EX2 5DW, UK; Analytic and Translational Genetics Unit, Massachusetts General Hospital, Boston, Massachusetts 02114 USA; Psychiatric & Neurodevelopmental Genetics Unit, Massachusetts General Hospital, Boston, Massachusetts, 02114, USA; Estonian Genome Center, Institute of genomics, University of Tartu, Tartu 51010, Estonia; Department of Obstetrics and Gynecology, Institute of Clinical Medicine, University of Tartu, Tartu, Estonia; The Wellcome Trust Centre for Human Genetics, Nuffield Department of Medicine, University of Oxford, Oxford, OX3 7BN, UK; Li Ka Shing Centre for Health Information and Discovery, Big Data Institute, University of Oxford, Oxford, UK; Genetics of Complex Traits, University of Exeter Medical School, University of Exeter, Exeter, EX2 5DW, UK; National Institute for Health Research Oxford Biomedical Research Centre, Oxford University Hospital, Old Road, Oxford OX3 7LE, UK; Clinical Trial Service Unit & Epidemiological Studies Unit (CTSU), Nuffield Department of Population Health, Big Data Institute Building, Roosevelt Drive, University of Oxford, Oxford, OX3 7LF, UK; Medical Research Council Population Health Research Unit at the University of Oxford, Nuffield Department of Population Health, University of Oxford, UK

## Abstract

GWAS of erectile dysfunction (ED) in 6,175 cases among 223,805 European men identified one new locus at 6q16.3 (lead variant rs57989773, OR 1.20 per C-allele; p = 5.71×10^−14^), located between *MCHR2* and *SIM1*. In-silico analysis suggests *SIM1* to confer ED risk through hypothalamic dysregulation; Mendelian randomization indicates genetic risk of type 2 diabetes causes ED. Our findings provide novel insights into the biological underpinnings of ED.

---

Erectile dysfunction (ED) is the inability to develop or maintain a penile erection adequate for sexual intercourse^1^. ED has an age-dependent prevalence, with 20-40% men aged 60-69 years affected^1^. The genetic architecture of ED remains poorly understood, owing in part to a paucity of well-powered genetic association studies.

We conducted a genome-wide association study (GWAS) using data from the population-based UK Biobank (UKBB) and the Estonian Genome Center of the University of Tartu (EGCUT) cohorts and hospital-recruited Partners HealthCare Biobank (PHB) cohort (Supplementary Methods).

The prevalence of ED (defined as self-reported, or physician-reported ED using ICD10 codes N48.4 and F52.2, or use of oral ED medication (sildenafil/Viagra, tadalafil/Cialis or vardenafil/Levitra), or a history of surgical intervention for ED (using OPCS-4 codes: L97.1 and N32.6)) in the cohorts was 1.53% (3,050/199,352) in UKBB, 7.04% (1,182/16,787) in EGCUT and 25.35% (1,943/7,666) in PHB (Supplementary Table 1).

GWAS in UKBB revealed a single genome-wide significant (p < 5×10^−8^) locus at 6q16.3 (Figures 1A and 1B; lead variant, rs57989773, EAFUKBB (C-allele) = 0.24; OR 1.23; p = 3.0×10^−11^). Meta-analysis with estimates from PHB (OR 1.20; p = 9.84×10^−5^) and EGCUT (OR 1.08; p = 0.16) yielded a pooled meta-analysis OR 1.20; p = 5.72×10^−14^ (Figure 1C). Meta-analysis of all variants yielded no further genome-wide loci. Meta-analysis of our results with previously suggested ED-associated variants did not result in any further significant loci (Supplementary Methods; Supplementary Table 2).

**FIGURE 1.**
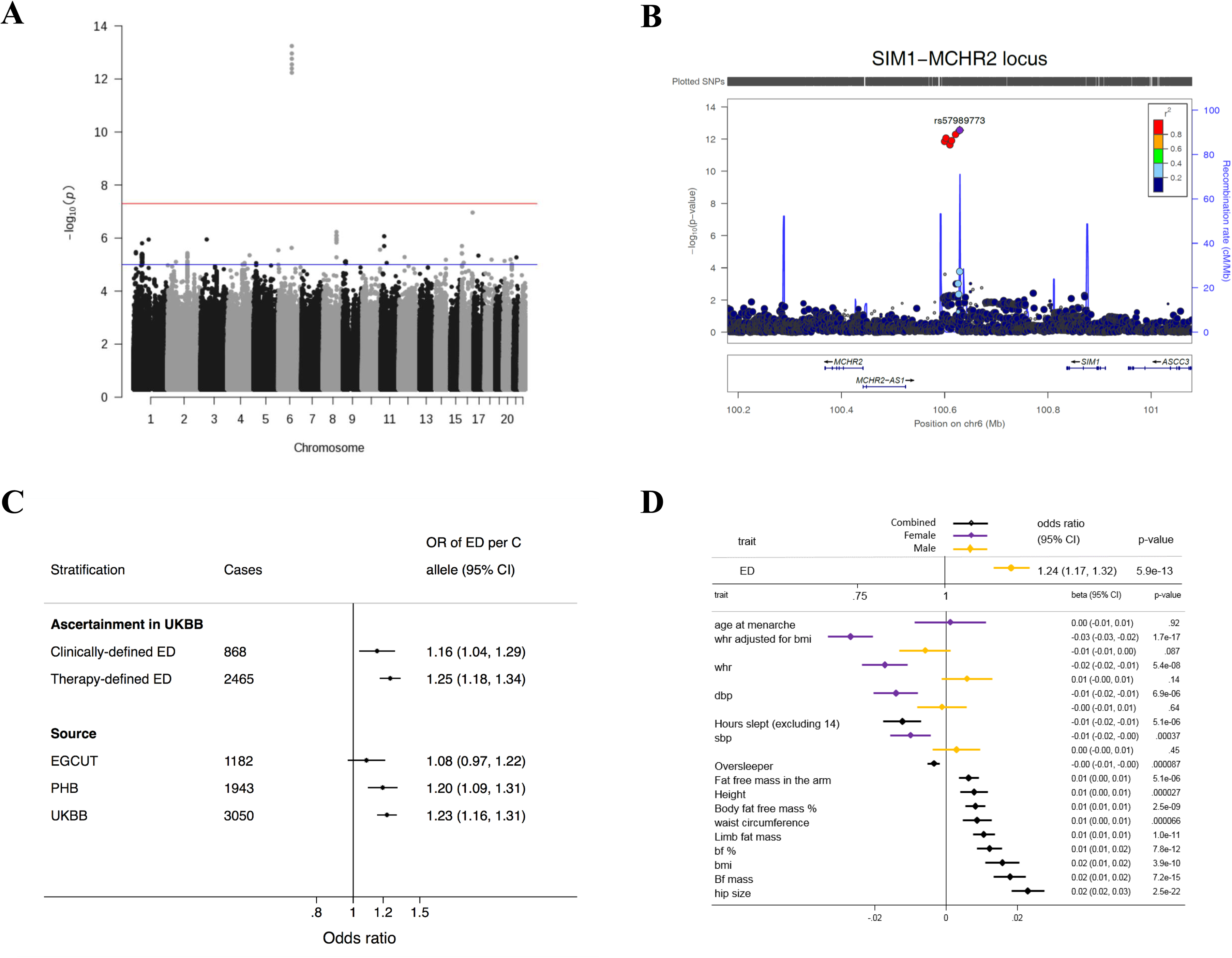
6q16.3 (LEAD VARIANT rs57989773) IS A NOVEL ED-ASSOCIATED LOCUS AND EXHIBITS PLEIOTROPIC PHENOTYPIC EFFECTS. A: **Genome-wide meta-analysis revealed a single genome-wide significant locus for ED at 6q16.3**. B: **Six genome-wide significant variants at 6q16.3 are in high LD**. C: **The association of rs57989773 with ED shows a consistent direction of effect across the three cohorts and across clinically- and therapy-defined ED in UKBB**. D: **PheWAS reveals sex-specific associations of rs57989773 with waist-hip ratio and blood pressure**. A PheWAS of 105 predefined traits using the lead ED SNP rs57989773 found associations with 12 phenotypes at p-value < 4.8 × 10^−4^ (surpassing the Bonferroni-corrected threshold of 0.05/105; Supplementary Table 3). Due to the nature of the ED phenotype and previously reported sex-specific effects in the *MCHR2-SIM1* locus, sex-specific analyses were performed in significant traits. Diastolic blood pressure (dbp) and systolic blood pressure (sbp) are included here (despite not meeting the Bonferroni-corrected threshold in the original analysis), due to previous reports of effects on blood-pressure in patients with rare, coding variants in *SIM1* and because the female-specific effects on blood pressure did meet the original threshold. Sexual heterogeneity was found to be significant (surpassing a Bonferroni-corrected threshold of 0.05/7 for the number of traits where sex-specific analyses were conducted) for diastolic blood pressure (p-value_heterogeneity_ = 6.52 × 10^−3^), systolic blood pressure (p-value_heterogeneity_ = 3.73 × 10^−3^), waist to hip ratio (p-value_heterogeneity_ = 2.39 × 10^−6^) and waist to hip ratio adjusted for BMI (p-value_heterogeneity_ = 1.77 × 10^−5^). Continuous traits were standardised prior to analysis to facilitate comparison.

The association of rs57989773 was consistent across clinically- and therapy-defined ED and across different ED drug classes (Figure 1C and Supplementary Figure S1). No further genome-wide significant loci were identified for ED when limited to clinically- or therapy-defined cases (Supplementary Notes).

A PheWAS of 105 predefined traits (Supplementary Table 3) using the lead ED SNP rs57989773 found associations with 12 phenotypes at p-value < 5×10^−4^ (surpassing the Bonferroni-corrected threshold of 0.05/105), including adiposity (9 traits), adult height and sleep-related traits. Sex-stratified analyses revealed sexual dimorphism for waist-hip ratio (WHR), systolic and diastolic blood pressure (Figure 1D and Supplementary Table 4).

rs57989773, the lead variant at the 6q16.3 locus, lies in the intergenic region between *MCHR2* and *SIM1*, with *MCHR2* being the closest gene (distances to transcription start sites of 187kb for *MCHR2* and 284kb for *SIM1*). Previous work has implicated the *MCHR2-SIM1* locus in sex-specific associations on age at voice-breaking and menarche^2^. The puberty timing-associated SNP in the *MCHR2-SIM1* region (rs9321659) was not in LD with our lead variant (r^2^=0.003) and was not associated with ED (p = 0.32) in our meta-analysis, suggesting that the ED locus represents an independent signal.

To identify the tissue and cell types in which the causal variant(s) for ED may function, we examined chromatin states across 127 cell types^3,4^ for the lead variant rs57989773 and its proxies (r^2^>0.8, determined using HaploReg v4.1 (Supplementary Methods)). Enhancer marks in several tissues, including embryonic stem cells, mesenchymal stem cells and endothelial cells, indicated that the ED-associated interval lies within a regulatory locus (Figure 2A, Supplementary Table 5).

**FIGURE 2.**
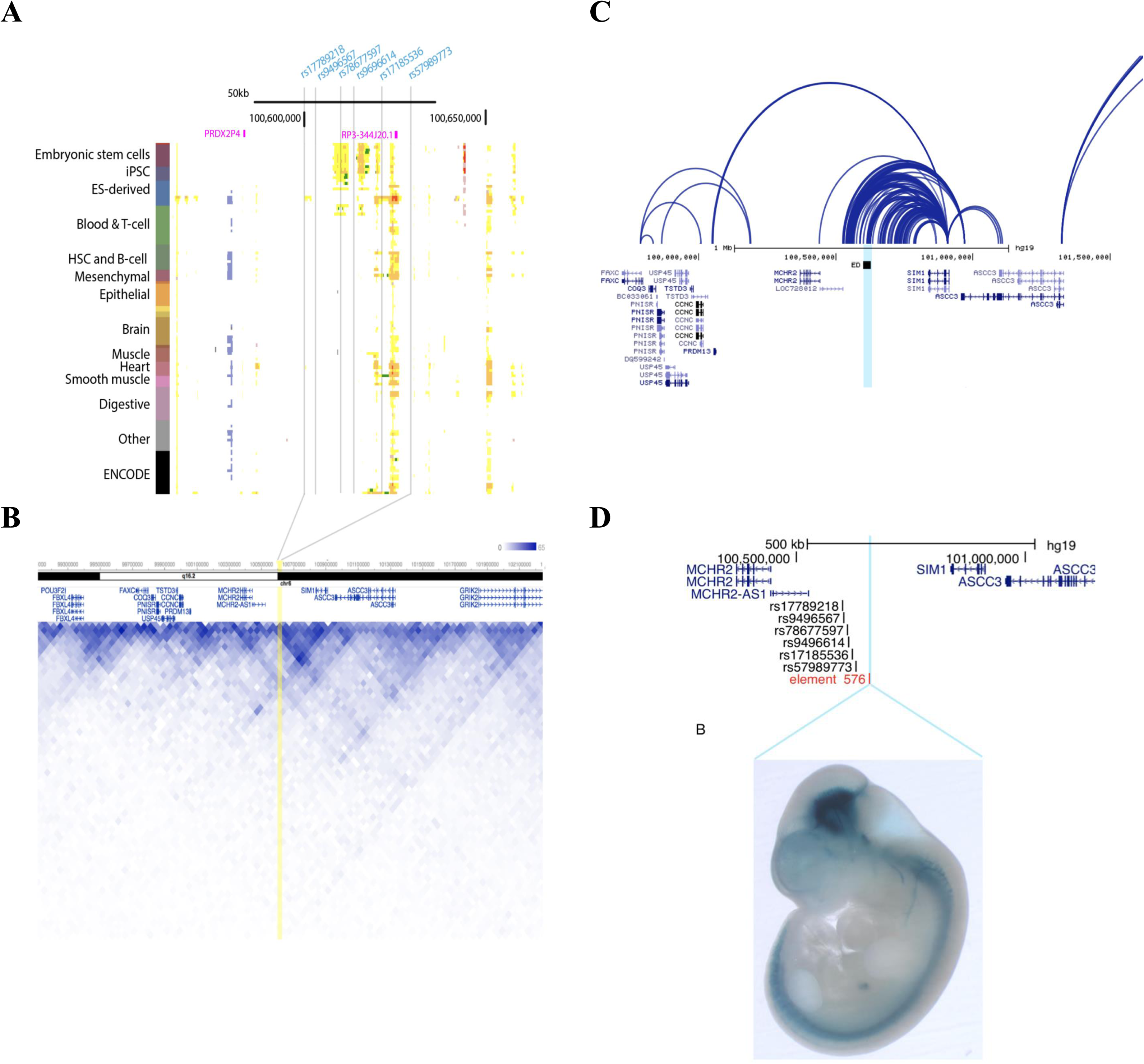
FUNCTIONAL ANALYSIS OF 6q16.3 IMPLICATES *SIM1* IN ED PATHOGENESIS. A: **Epigenomic signals surrounding the ED-associated region**. Chromatin state annotations for the ED-associated region across 127 reference epigenomes (rows) for cell and tissue types profiled by the Roadmap Epigenomics Project^1,2^. Blue vertical lines indicate the position of the ED-associated variant (rs57989773) and its proxies that are in LD r2>0.8 determined using HaploReg v4.1^3^ (rs17789218, rs9496567, rs78677597, rs9496614, and rs17185536). The purple block labelled ‘rp3-344J20.1’ represents the arginase 2 processed pseudogene (LOC100129854). B: **The ED-associated interval is functionally connected to the SIM1 promoter in embryonic stem cells**. The 3D Genome Browser^4^ was used to visualize chromosome conformation capture (Hi-C) interactions contact probabilities in human embryonic stem cells^5^, revealing high contact probability between the ED-associated region (highlighted in yellow) and *SIM1* at 40-kb resolution. The yellow vertical line represents the location of the ED-associated interval. The heat map values on a color scale correspond to the number of times that reads in two 40-kb bins were sequences together (bluestronger interaction, whitelittle or no interaction). C: **The *MCHR2-SIM1* intergenic region forms functional connections to the *SIM1* promoter in endothelial progenitors**. The 3D Genome Browser^4^ was used to visualize Capture Hi-C in endothelial precursors (Data from Fraser lab). Light blue vertical line indicates position of the ED-associated interval. D: **The *MCHR2-SIM1* intergenic region harbors a neuronal enhancer**. Upper panel: Position of human element hs576 (blue vertical line) and the ED-associated variant rs57989773 and its 5 proxies in r2>0.8 (rs17789218, rs9496567, rs78677597, rs9496614, rs17185536). hs576 is flanked by genes *MCHR2-AS1SIM1*. This panel was generated using the UCSC genome browser^6^. Lower panel: Expression pattern of human element hs576 in a mouse embryo at e11.5. Expression pattern shows that hs576 drives *in vivo* enhancer activity specifically in mesencephalon (midbrain) and cranial nerve. Expression data were derived from the VISTA enhancer browser^7^.

To predict putative targets and causal transcripts, we assessed domains of long-range three-dimensional chromatin interactions surrounding the ED-associated interval (Figure 2B). Chromosome conformation capture (Hi-C) in human embryonic stem cells^5^ showed that *MCHR2* and *SIM1* were in the same topologically associated domain (TAD) as the ED-associated variants, with high contact probabilities (referring to the relative number of times that reads in two 40-kb bins were sequenced together) between the ED-associated interval and *SIM1* (Figure 2B and Figure S2).

This was further confirmed in endothelial precursor cells^6^, where Capture Hi-C revealed strong connections between the *MCHR2-SIM1* intergenic region and the *SIM1* promoter (Figure 2C), pointing towards *SIM1* as a likely causal gene at this locus.

We next used the VISTA enhancer browser^7^ to examine *in vivo* expression data for non-coding elements within the *MCHR2-SIM1* locus. A regulatory human element (hs576), located 30-kb downstream of the ED-associated interval, seems to drive *in vivo* enhancer activity specifically in the midbrain (mesencephalon) and cranial nerve in mouse embryos (Figure 2D). This long-range enhancer close to ED-associated variants recapitulated aspects of *SIM1* expression (Figure 2D), further suggesting that the ED-associated interval belongs to the regulatory landscape of *SIM1*. Taken together these data suggest that the *MCHR2-SIM1* intergenic region harbors a neuronal enhancer and that *SIM1* is functionally connected to the ED-associated region.

Single-minded homolog 1 (*SIM1*) encodes a transcription factor that is highly expressed in hypothalamic neurons^8^. Rare variants in *SIM1* have been linked to a phenotype of severe obesity and autonomic dysfunction^9,10^, including lower blood pressure. A summary of the variant-phenotype associations at the 6q16 locus in human and rodent models is shown in Supplementary Table 6. Post-hoc analysis of association of rs57989773 with autonomic traits showed nominal association with syncope, orthostatic hypotension and urinary incontinence (Figure S3). The effects on blood pressure and adiposity seen in patients with rare coding variants in *SIM1* are recapitulated in individuals harbouring the common ED-risk variant at the 6q16.3 locus (Figure 1D, Supplementary Figure S3), suggesting that *SIM1* is the causal gene at the ED-risk locus. *Sim1*-expressing neurons also play an important role in the central regulation of male sexual behavior as mice that lack the melanocortin receptor 4 (*MC4R*) specifically in *Sim1*-expressing neurons show impaired sexual performance on mounting, intromission, and ejaculation^11^. Thus, hypothalamic dysregulation of *SIM1* could present a potential mechanism for the effect of the *MCHR2-SIM1* locus on ED.

An additional functional mechanism may be explained by proximity of the lead variant (rs57989773) to an arginase 2 processed pseudogene (LOC100129854), a long non-coding RNA (Figure 2A). RPISeq^12^ predicts that the pseudogene transcript would interact with the ARG2 protein, with probabilities of 0.70-0.77. Arginine 2 is involved in nitric oxide production and has a previously established role in erectile dysfunction^13,14^. GTEX expression data^15^ demonstrated highest mean expression in adipose tissue, with detectable levels in testis, fibroblasts and brain. Expression was relatively low in all tissues however, and there was no evidence that any SNPs associated with the top ED signal were eQTLs for the *ARG2* pseudogene or *ARG2* itself.

As a complementary approach, we also used the Data-driven Expression Prioritized Integration for Complex Traits and GWAS Analysis of Regulatory or Functional Information Enrichment with LD correction (DEPICT and GARFIELD respectively; Supplementary Methods)^16,17^ tools to identify gene-set, tissue-type and functional enrichments. In DEPICT, the top two prioritized gene-sets were ‘regulation of cellular component size’ and ‘regulation of protein polymerization’, whereas the top two associated tissue/cell types were ‘cartilage’ and ‘mesenchymal stem cells’. None of the DEPICT enrichments reached an FDR threshold of 5% (Supplementary Tables 7-9). GARFIELD analyses also did not yield any statistically significant enrichments.

LD score regression^18,19^ identified ED to be correlated and share genetic architecture with type 2 diabetes (rg = 0.40, nominal p-value = 0.0008; FDR-adjusted p-value = 0.0768; Supplementary Table 10). Mendelian randomization^20^ (Supplementary Tables 11-17) identified genetic risk to T2D to be causally implicated in ED: OR 1.11 (95% CI 1.05-1.17, p = 3.5×10^−4^, per 1-log higher genetic risk of T2D; with insulin resistance likely representing a mediating pathway. A potential causal effect of SBP was also identified, with higher SBP being linked to higher risk of ED. In keeping with this, genetic risk of CHD showed weak effects on risk of ED, suggesting that pathways leading to CHD may be implicated in ED.

In contrast, no causal effects of BMI (using a polygenic score or a single SNP in *FTO*) or education on ED were identified. This suggests the effect of the rs57989773 on ED is independent of its effect on BMI.

We also looked at variation at the 4q26 locus, containing *PDE5A* which encodes phosphodiesterase 5 (PDE5) - the primary drug target for PDE5-inhibitors such as sildenafil. Of all 4,670 variants within a 1Mb window of *PDE5A* (chromosome 4:119,915,550-121,050,146 as per GRCh37/hg19), the variant with the strongest association was rs115571325, 26Kb upstream from *PDE5A* (OR_Meta_ 1.25, nominal p-value = 8.46 x 10^−4^; Bonferroni-corrected threshold (0.05/4,670) = 1.07 x 10^−5^; Figure S4).

In conclusion, our GWAS of 6,175 ED cases, the largest to date, identifies a new locus associated with ED, and provides evidence implicating an effect of common non-coding variants on *SIM1*. We also show genetic risk to T2D as causally implicated in the aetiology of ED, with suggestive evidence for blood pressure and coronary heart disease. Further large-scale GWAS of ED are needed in order to provide additional clarity on its genetic architecture, aetiology and shed light on potential new therapies.

## Acknowledgements

We thank the UK Biobank (http://www.ukbiobank.ac.uk/; application 11867), Partners HealthCare Biobank https://biobank.partners.org/), and the Estonian Biobank of the Estonian Genome Center of the University of Tartu (https://www.geenivaramu.ee/en) and their participants.

---

**Figure S1.**
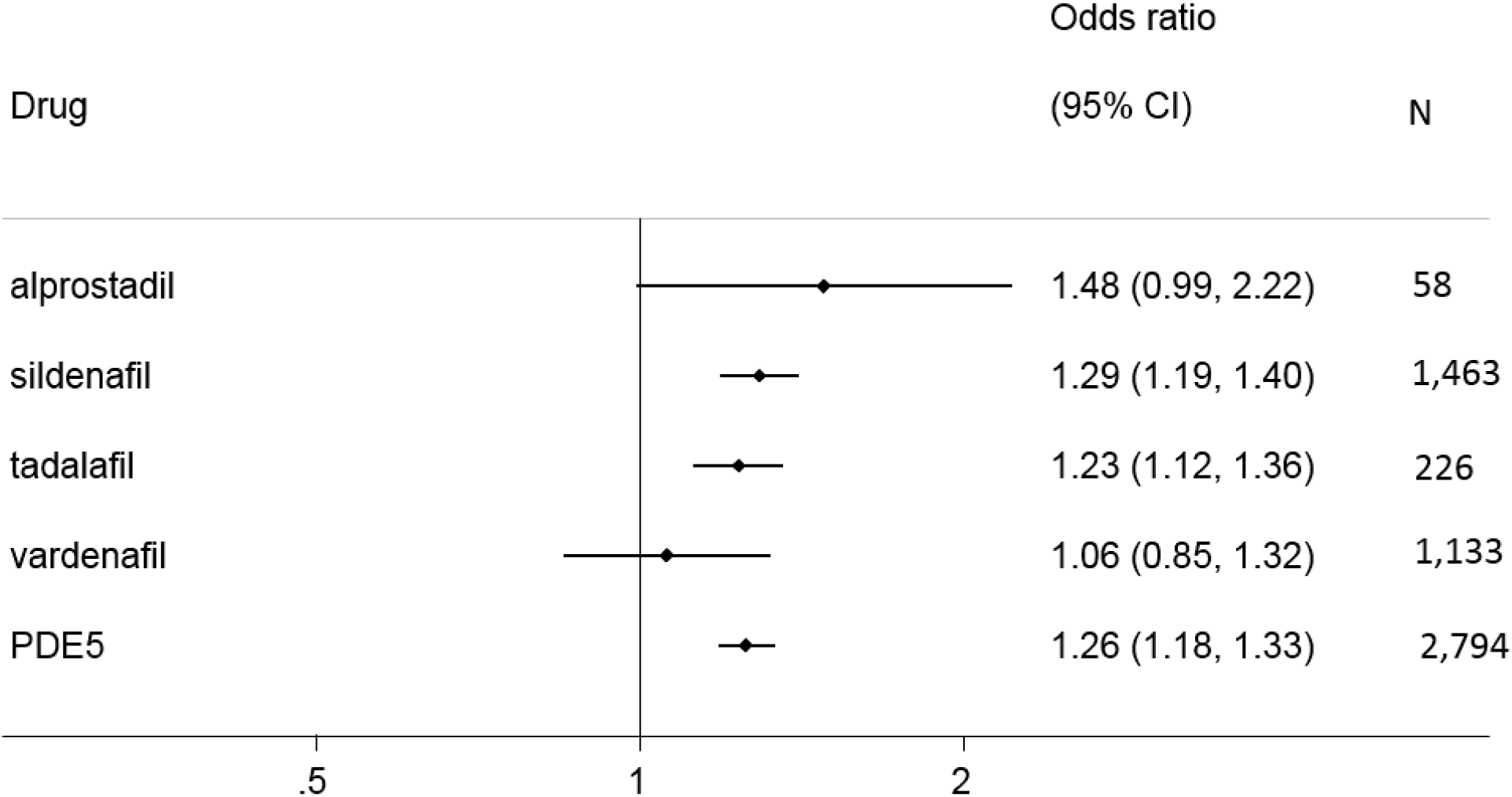
The association of rs57989773 remains consistent across different ED drug classes.

**Figure S2.**
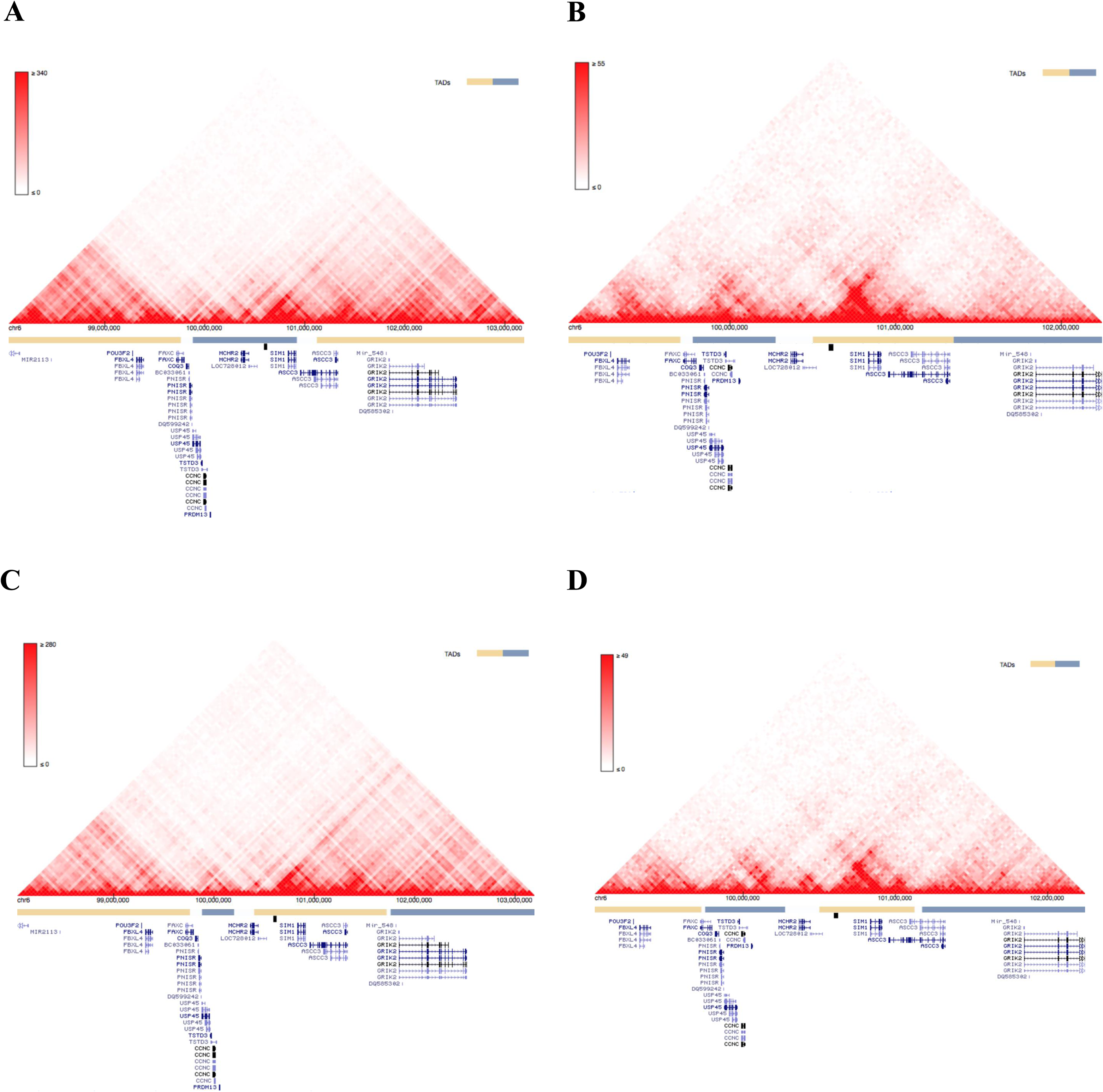
Hi-C interaction maps in several cell types. The 3D Genome Browser^4^ was used to visualize the spatial organisation surrounding the ED-associated region. Heatmap shows chromosome conformation capture (Hi-C) interactions contact probabilities in (A) human MES mesendoderm cells^5^ at 40-kb resolution; (B) human endothelial progenitors (HUVEC) at 25-Kb resolution^8^; (C) human mesenchymal stem cells (MSC)^5^ at 40-kb resolution; and (D) human endothelial progenitors (HUVEC) at 25-kb^8^ resolution. The heat map values on a colour scale correspond to the number of times that reads in two 40-kb bins were sequences together (red - stronger interaction, white - little or no interaction). The second panel indicates the location of the ED-associated region. The third panel shows the UCSC reference genes

**Figure S3.**
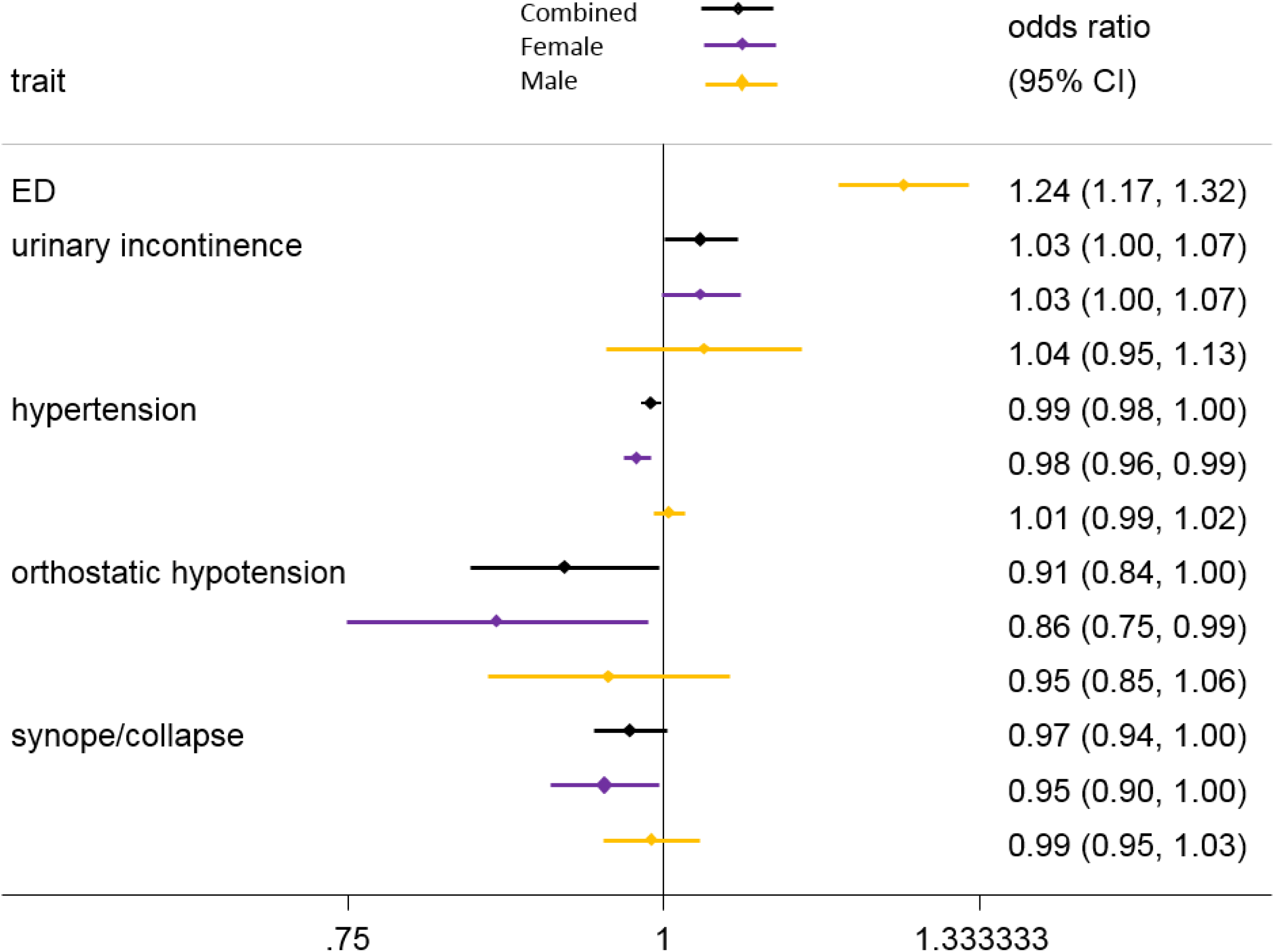
Association of rs57989773 with autonomic phenotypes.

**Figure S4.**
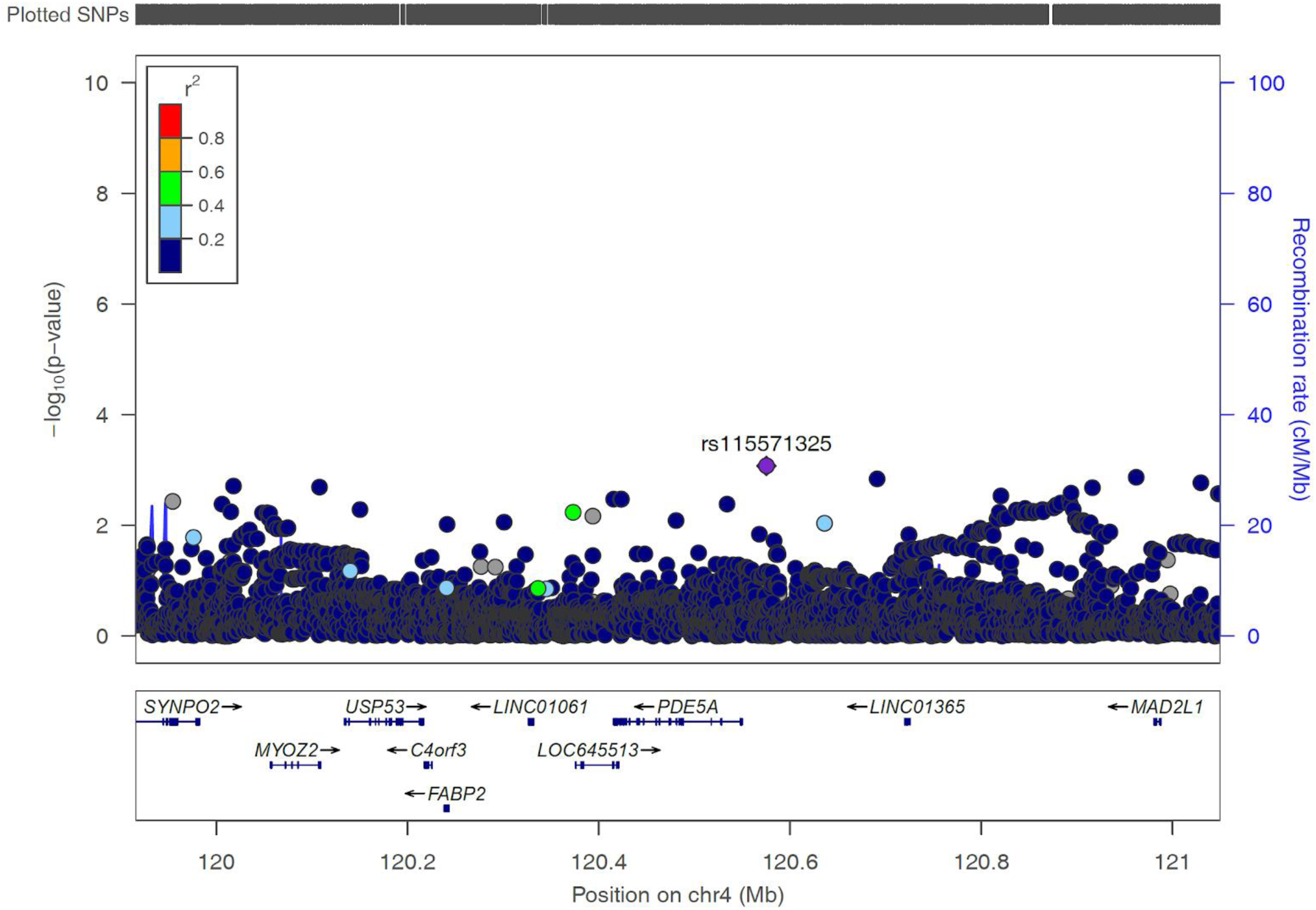
Association of variants in PDE5A region with ED.

