## Supplementary Materials for "GWAS identifies novel risk locus for erectile dysfunction and implicates hypothalamic neurobiology and diabetes in etiology"

STUDY SUBJECTS 2
Study subjects: Partners HealthCare Biobank 2
Study subjects: UK Biobank 2
Study subjects: Estonian Genome Center of the University of Tartu 3
GENOTYPING, QC, AND IMPUTATION 4
Genotyping, QC, and imputation: Partners Healthcare Biobank 4
Genotyping, QC, and imputation: UK Biobank 4
Genotyping, QC, and imputation: Estonian Genome Center of the University of Tartu 5
STATISTICAL ANALYSES 6
Statistical analyses: Partners Healthcare 6
Statistical analyses: UK Biobank 6
Statistical analyses: Estonian Genome Center of the University of Tartu 6
Statistical analyses: Meta-analyses 6
Replication and meta-analysis of previously reported ED-associated SNPs 7
IN-SILICO FUNCTIONAL FOLLOW-UP 8
Phenome-wide Association Scan (PheWAS) 8
DEPICT 8
GARFIELD 9
LD Score regression and cross-trait genetic correlation analysis 9
Mendelian Randomization 9
RESOURCES/ URLs 11
ACKNOWLEDGEMENTS 12
REFERENCES 13

### SUPPLEMENTARY METHODS

#### STUDY SUBJECTS

##### Study subjects: Partners HealthCare Biobank

We identified cases of erectile dysfunction (ED) and healthy male controls from the Partners HealthCare Biobank[1,2], a biorepository of consented patient samples at Partners HealthCare (the parent organization of Massachusetts General Hospital and Brigham and Women's Hospital). All patients who participate in the Partners Biobank are consented for their samples to be linked to their identified clinical information.

The ED cases (Supplementary Table 1) were identified by querying the Partners Biobank with ICD-10 code N52 (male erectile dysfunction). In addition to ICD-10 code, we also identified ED cases by querying the Partners Biobank with the following drug prescriptions: sildenafil, viagra (25mg, 50mg, and 100mg tablet), tadalafil, cialis (10mg and 20mg tablet), vardenafil, levitra (5mg, 10mg, and 20mg tablet). Patients with pulmonary hypertension were excluded from the identified ED case pool. The controls were males from the Partners Biobank. We also extracted age for ED cases and controls from the Biobank for subsequent analyses. There are 1,943 ED cases and 5,723 male controls with genome-wide genotyped data.

##### Study subjects: UK Biobank

UK Biobank (UKBB) is a prospective study of more than 500,000 British individuals recruited from 2006 to 2010, aged between 45 and 69[3]. Phenotypic information available includes self-reported medical history (including medication use) as ascertained by verbal interview at enrollment and hospital-derived electronic health record (EHR) data, including International Classification of Disease (ICD-10) diagnosis codes and Office of Population and Censuses Surveys (OPCS-4) procedure codes.

Individuals in UKBB were defined as having ED (Supplementary Table 1) on the basis of at least one of the following criteria: Self-reported ED/impotence at time of enrolment; hospitalisation for ICD-10 codes N48.4, F52.2 or N52; hospitalisation for OPCS-4 coded procedures L97.1 or N32.6; or self-reported ED medication (either sildenafil, Viagra, tadalafil, Cialis, vardenafil, Levitra) use.

Of the 488,377 individuals with available genotype data (see section **Genotyping, QC, and imputation: UK Biobank** below), we excluded individuals that had: withdrawn their consent for participation; high sample heterozygosity and missingness; >10 third degree relatives; putative sex chromosome aneuploidy; sex mismatches (genetic vs. self-reported and between assessments); ethnicity mismatches (genetic vs self-reported for White British individuals, and mismatches between assessments) and diagnostic codes for pulmonary hypertension. After applying these filters, 199,352 male subjects remained, of whom 3,050 met the case-definition criteria for ED, whereas 196,302 subjects served as controls.

**Study subjects: Estonian Genome Center of the University of Tartu**

The Estonian Genome Center of the University of Tartu (EGCUT) is a population-based biobank with a current cohort size of 51,515 participants[4]. Upon recruitment, the biobank participants have filled out a thorough questionnaire, covering lifestyle, diet and clinical diagnoses (described by ICD-10 codes). Data are periodically updated by linking with national health registries. In EGCUT, ED cases (Supplementary Table 1) were defined using ICD-10 codes F52.2 or N48.4; or data on prescribed drugs (with active compounds tadalafil, sildenafil, vardenafil). Males without any of these diagnosis codes or prescribed drugs were used as controls. The analysis included a total of 1,182 male cases and 15,605 male controls.

### GENOTYPING, QC, AND IMPUTATION

##### Genotyping, QC, and imputation: Partners Healthcare Biobank

DNA samples from the patients in the Partners Biobank were extracted from whole blood. A total of 20,087 samples were genotyped with Illumina Multi-Ethnic Genotyping Array (first batch), Expanded Multi-Ethnic Genotyping Array (second batch), and Multi-Ethnic Global BeadChip (third batch), all of which were designed to capture the global diversity of genetic backgrounds. The number of genotyped variants ranged from 1,416,020 to 1,778,953. We performed QC on each genotyping batch separately as follows: we removed single nucleotide polymorphisms (SNPs) with genotype missing rate > 0.05 before sample-based QC; excluded samples with genotype missing rate > 0.02, absolute value of heterozygosity > 0.2, or failed sex checks; removed SNPs with missing rate > 0.02 after sample-based QC. To merge genotyping batches for imputation and analyses, we performed batch QC by removing SNPs with significant batch association (p-value < $1.0x10^{-6}$ between different batches). Since the Partners Biobank samples have diverse population backgrounds, we performed Hardy-Weinberg equilibrium test (p-value < $1.0x10^{-6}$) for SNP-based QC after extracting samples with European ancestry (see below). We also performed relatedness tests by identifying pairs of samples with $\pi > 0.2$ and excluding one sample from each related sample pair (560 samples excluded). All QC were conducted using PLINK v1.9 and R software.

We extracted samples with European ancestry based on principal component analysis (PCA) with 1000 Genomes Project reference samples. Details of the procedure used to extract European ancestry samples were described previously [5,6]. Briefly, we ran PCA on study samples combined with 1000 Genomes Project reference samples and calculate Euclidean distance ($d_{EUR}$) for each study sample to the average PC1 and PC2 of the 1000 Genomes Project EUR samples. A total of 16,453 study samples with European ancestry were extracted based on $d_{EUR} < 0.003$. We then performed PCA on the European ancestry samples to obtain PCs for the subsequent analyses.

Genotype imputation was performed on the QCed European ancestry samples with a 2-step pre-phasing/imputation approach. We used Eagle2 for the pre-phasing and minimac3 for imputation, with a reference panel from 1000 Genomes Project phase 3. The final analytic data includes 1,943 ED cases and 5,723 controls of European ancestry with imputed genotype data.

##### Genotyping, QC, and imputation: UK Biobank

Genotyping, quality control and imputation were performed centrally by UKBB, and details are described elsewhere[7] (see also http://biobank.ctsu.ox.ac.uk/crystal/label.cgi?id=100319). Briefly, genotype data is available for 488,377 individuals, 49,950 of whom were genotyped using the Applied Biosystems™ UK BiLEVE Axiom™ Array by Affymetrix (containing 807,411 markers[8]). The remaining 438,427 individuals were genotyped using the Applied Biosystems™ UK Biobank Axiom™ Array by Affymetrix (containing 825,927 markers). Both of these arrays were specifically designed for use in the UKBB project and share ~95% of marker content. Phasing was done using SHAPEIT3, and imputation was conducted using IMPUTE4. For imputation, the Haplotype Reference Consortium (HRC) panel[9] was used wherever possible, and for SNPs not in that reference panel, a merged UK10K + 1000 Genomes reference panel was used. SNPs were imputed from both panels, but the HRC imputation was preferentially used for SNPs present in both panels. Given known issues with non-HRC imputed SNPs with the current UKBB genotype data release (http://www.ukbiobank.ac.uk/2017/07/important-note-about-imputed-genetics-data/), we included only HRC-imputed SNPs in our dataset.

**Genotyping, QC, and imputation: Estonian Genome Center of the University of Tartu**

In EGCUT, DNA was extracted from whole blood. Genotyping was carried out using Illumina Human CoreExome, OmniExpress, 370CNV BeadChip and GSA arrays. Genotype array data was filtered sample-wise by excluding on the basis of call rate (<98%), heterozygosity (>mean±3 SD), genotype and phenotype sex discordance, cryptic relatedness (IBD>20%) and outliers from the European descent based on the MDS plot in comparison with HapMap reference samples. SNP quality filtering included call rate (<99%), MAF (<1%) and extreme deviation from Hardy–Weinberg equilibrium (P-value$<1 \times 10^{-4}$). Imputation was performed on the QCed samples, using SHAPEIT2 for prephasing, the Estonian-specific reference panel[10] and IMPUTE2 with default parameters.

### STATISTICAL ANALYSES

##### Statistical analyses: Partners Healthcare

Logistic regression was used to test genome-wide association for ED in PHB, adjusted for 10 PCs and age, using PLINK v1.9[11].

##### Statistical analyses: UK Biobank

We used BOLT-LMM[12] v2.3 to perform association analyses in UKBB. BOLT-LMM computes statistics for testing association between phenotype and genotypes using a linear mixed model (LMM)[12]. The performance of BOLT-LMM on the UKBB dataset has been previously validated, allows for inclusion of related individuals and has been shown to significantly increase power when compared to traditional linear regression methods[13]. All analyses were adjusted for age.

SNPTEST[14] v2.5.2 was used to compute confirm association statistics for variants with a p-value of association of less than $5 \times 10^{-8}$ on the BOLT-LMM analyses.

##### Statistical analyses: Estonian Genome Center of the University of Tartu

EPACTS v3.3.0 (using option q.emmax) was used to perform association testing in EGCUT, adjusting for age at recruitment and the kinship matrix.

##### Statistical analyses: Meta-analyses

Prior to meta-analysis, we performed standardized study-level quality control using EASYQC[15]. All three studies (UKBB, PHB, and EGCUT) were subjected to the same QC measures, with inclusion of variants with imputation INFO scores > 0.4 and MAF > 1%. Indels and CNVs were not included in the meta-analysis. Genomic control correction was applied to each dataset prior to meta-analysis.

METAL software[16] was used for performing fixed effects inverse-variance weighted meta-analysis of allelic effect sizes after conversion onto the log-odds scale. This has been shown to be a valid method for meta-analyses of GWA studies of binary phenotypes using linear mixed models[17].

In addition to the main meta-analysis, we conducted two additional meta-analyses using clinically- or therapy-defined cases respectively. Since only UKBB and PHB had this data available, only these two cohorts were included in these analyses. Identical methodology as described above was followed when performing these meta-analyses.

**Replication and meta-analysis of previously reported ED-associated SNPs**

We performed a literature review for previous ED GWAS and identified three studies[18–20]. We extracted all independent, autosomal SNPs associated with ED with $p \leq 9 \times 10^{-6}$ (i.e. all autosomal SNPs included on the GWAS Catalog database entry for "impotence" or "erectile dysfunction"), yielding 23 SNPs. Summary statistics for these variants were subsequently extracted from our GWAMA results. We then performed effective sample-size weighted Z-score meta-analyses of previously reported summary statistics for 17 variants (3 variants omitted due to MAF < 1% in all three cohorts in our study, 2 variants omitted due to absence of effect size direction in the original report and 1 due to alleles being inconsistent between studies) with summary statistics extracted from our GWAMA (Supplementary Table 2).

#### IN-SILICO FUNCTIONAL FOLLOW-UP

UCSC Genome Browser (available at http://genome.ucsc.edu/; December 2013 (GRCh37/hg19) assembly) was used to determine distance of rs57989773 to the transcription start sites of MCHR2 and SIM1.

##### Phenome-wide Association Scan (PheWAS)

A PheWAS of traits in UKBB was carried out as previously described[21]. Briefly, a range of phenotypes were available in UK Biobank, derived from self-reported questionnaire data, ICD10 diagnoses and baseline measurements at clinic visits as part of the study. We tested the association of the lead ED SNP rs57989773 with a range of phenotypes and traits including: anthropometric, reproductive, cardiovascular, learning/memory and incidence of various diseases (Supplementary Table 3). Association testing was carried out with inverse normalised phenotypes to account for any skewed distributions, using linear regression models in STATA 13, adjusting for SNP chip type (UKB Axiom or UK BiLEVE), ancestry-principal components 1 to 5 supplied by UK Biobank, test centre and age (or year of birth for age at menarche) with the exception of three traits: hypertension, hypothyroidism and household income. Hypertension and hypothyroidism were tested using logistic regression and household income was tested using ordinal logistic regression. The logistic and ordinal logistic models were adjusted using the same covariates as the linear regression model. Due to the nature of the ED phenotype and previously reported sex-specific effects in the MCHR2-SIM1 locus, sex-specific analyses was performed on significant traits. Testing for heterogeneity was performed to assess whether any observed sex-specific effects were significant after accounting for multiple testing.

##### DEPICT

DEPICT[22] (Data-driven Expression Prioritized Integration for Complex Traits) is a comprehensive pathway analysis tool. We used DEPICT to prioritise likely causal genes at associated loci, and to identify enriched gene sets and tissue and cells types where genes from prioritised loci are highly expressed.

In our study, independent variants were identified in the genome-wide association meta-analysis result using PLINK v.1.9 to clump SNPs at an LD-threshold of $r^2$=0.1 and a physical distance threshold of 500kb, resulting in 37 independent variants with a p-value threshold of $p < 1 \times 10^{-5}$. SNPs in HLA regions, on sex chromosomes or not present in 1000 Genomes Project were excluded from DEPICT analysis. DEPICT was used to identify tissue and cell type annotations in which genes from associated regions were highly expressed, to identify reconstituted gene sets enriched for genes from associated regions and to prioritize genes within associated regions.

**GARFIELD**

GARFIELD is a functional enrichment analysis approach described more fully elsewhere[23]. Briefly, GARFIELD is a nonparametric method to assess enrichment of GWAS signals in regulatory of functional regions in different cell-types. The software LD-prunes the GWAS-data before assessing fold enrichment at different p-value thresholds from the GWAS study of interest. To minimize bias, it takes into account LD-structure, gene density, and allele frequencies. GARFIELD analyses did not yield any statistically significant enrichments.

**LD Score regression and cross-trait genetic correlation analysis**

LD Hub[24] was used to conduct LD Score regression and cross-trait genetic correlation analysis. LD Hub is a centralized database of summary-level GWAS results for >100 diseases/traits from different publicly available resources/consortia and uses web interface that automates the LD Score regression and cross-trait genetic correlation analysis pipeline.

LD Score regression[25] quantifies the contribution of true polygenic signal and confounding biases, such as cryptic relatedness and population stratification, to inflated distribution of test statistics in genome-wide association studies, by examining the relationship between test statistics and linkage disequilibrium (LD). The LD Score regression intercept can be used to estimate a more powerful and accurate correction factor than genomic control.

Genetic correlation analysis was conducting using cross-trait LD Score regression[26]. This is a technique estimating genetic correlation that requires only GWAS summary statistics and is not biased by sample overlap.

**Mendelian Randomization**

Multiple traits have shown association with ED in observational studies, including BMI, educational attainment, hypertension, diabetes, and cardiovascular disease[27]. We therefore investigated the causal effects of these traits using Mendelian randomization (MR) (Supplementary Table 11). All MR analyses were performed in R 3.4.3 (R Development Core Team (2008). R: A language and environment for statistical computing. R Foundation for Statistical Computing, Vienna, Austria. ISBN 3-900051-07-0, Available at http://www.R-project.org) using the *MendelianRandomization* package[28]. Instrument rsIDs were updated using Python version 3.5.2 (Python Software Foundation. Python Language Reference, version 3.5.2. Available at http://www.python.org), libraries Pandas[29] and Biopython[30], and the NCBI SNP website (Available from: https://www.ncbi.nlm.nih.gov/snp/). Proxies ($r^2$ >0.8) were identified using SNiPA[31]. Sensitivity analyses, including removal of palindromic SNPs, had no effect on the MR estimates.

#### RESOURCES/ URLs

PLINK v1.9
URL: www.cog-genomics.org/plink/1.9/

BOLT-LMM v2.3
URL: https://data.broadinstitute.org/alkesgroup/BOLT-LMM/

EPACTS v3.3.0
URL: https://github.com/statgen/EPACTS

EASYQC v9.2
URL: http://www.uni-regensburg.de/medizin/epidemiologie-praeventivmedizin/genetische-epidemiologie/software/

METAL
URL: http://csg.sph.umich.edu/abecasis/metal/

SNPTEST v2.5.2
URL: https://mathgen.stats.ox.ac.uk/genetics_software/snptest/snptest.html#introduction

LD HUB v1.9.0
URL: http://ldsc.broadinstitute.org/

DEPICT v1
URL: https://github.com/perslab/depict

MendelianRandomization v0.2.2 (R package)
URL: https://cran.r-project.org/web/packages/MendelianRandomization/index.html

HAPLOREG v4.1
URL: http://archive.broadinstitute.org/mammals/haploreg/haploreg.php

GARFIELD v2
URL: https://www.ebi.ac.uk/birney-srv/GARFIELD/

RPISeq v1.0
URL: http://pridb.gdcb.iastate.edu/RPISeq/references.php

#### ACKNOWLEDGEMENTS

##### LD Hub

We gratefully acknowledge all the studies and databases that made GWAS summary data available: ADIPOGen (Adiponectin genetics consortium), C4D (Coronary Artery Disease Genetics Consortium), CARDIoGRAM (Coronary ARtery DIsease Genome wide Replication and Meta-analysis), CKDGen (Chronic Kidney Disease Genetics consortium), dbGAP (database of Genotypes and Phenotypes), DIAGRAM (DIAbetes Genetics Replication And Meta-analysis), ENIGMA (Enhancing Neuro Imaging Genetics through Meta-Analysis), EAGLE (EArly Genetics & Lifecourse Epidemiology Eczema Consortium, excluding 23andMe), EGG (Early Growth Genetics Consortium), GABRIEL (A Multidisciplinary Study to Identify the Genetic and Environmental Causes of Asthma in the European Community), GCAN (Genetic Consortium for Anorexia Nervosa), GEFOS (GEnetic Factors for OSteoporosis Consortium), GIANT (Genetic Investigation of ANthropometric Traits), GIS (Genetics of Iron Status consortium), GLGC (Global Lipids Genetics Consortium), GPC (Genetics of Personality Consortium), GUGC (Global Urate and Gout consortium), HaemGen (haemotological and platelet traits genetics consortium), HRgene (Heart Rate consortium), IIBDGC (International Inflammatory Bowel Disease Genetics Consortium), ILCCO (International Lung Cancer Consortium), IMSGC (International Multiple Sclerosis Genetic Consortium), MAGIC (Meta-Analyses of Glucose and Insulin-related traits Consortium), MESA (Multi-Ethnic Study of Atherosclerosis), PGC (Psychiatric Genomics Consortium), Project MinE consortium, ReproGen (Reproductive Genetics Consortium), SSGAC (Social Science Genetics Association Consortium) and TAG (Tobacco and Genetics Consortium), TRICL (Transdisciplinary Research in Cancer of the Lung consortium), UK Biobank. We gratefully acknowledge the contributions of Alkes Price (the systemic lupus erythematosus GWAS and primary biliary cirrhosis GWAS) and Johannes Kettunen (lipids metabolites GWAS).
